# Gut microbiota promotes pain chronicity in Myosin1A deficient male mice

**DOI:** 10.1101/2023.02.21.529352

**Authors:** Ana Reynders, Z. Anissa Jhumka, Stéphane Gaillard, Guillaume Hoeffel, Annabelle Mantilleri, Pascale Malapert, Chiara Salio, Sophie Ugolini, Francis Castets, Andrew J. Saurin, Matteo Serino, Aziz Moqrich

## Abstract

Over the past decade, the gut microbiota has emerged as an important regulator of nervous system’s health and disease states^1^. Yet, its contribution to the pathogenesis of chronic somatic pain remains poorly documented. Chronic pain is a heavily debilitating disease affecting more than 1.5 billion people worldwide that can manifest through a long-lasting hypersensitivity to mechanical and/or thermal stimulations^2,3^. Maladaptive responses of dorsal root ganglia (DRG) neurons and spinal cord (SC) interneurons to tissue injuries and also of non-neuronal cells including DRG macrophages and SC microglia are acknowledged as important drivers of sensory symptoms underlying chronic pain^4,3,5–7^. Recent evidence shows that signals from gut microbiota are required for the initiation of injury-induced sensory hypersensitivity, via the ability to interact with the immune system^8–11^. However, whether and how gut microbiota promotes pain chronicity remains unknown. Here, we report that male mice lacking *Myosin1a* (KO)^12^ raised under single genotype housing conditions (KO-SGH) are predisposed to develop chronic injury-induced mechanical pain. We demonstrate that this predisposition is caused by their dysbiotic gut microbiota, which sustains the immune response in the DRG following neuropathic injury. Parental antibiotic treatment modifies gut microbiota composition and completely rescues the injury-induced chronic pain and associated DRG inflammatory response in male KO-SGH offspring. Together, our data establish a causal relationship between a dysbiotic gut microbiota and the predisposition to injury-induced chronic pain.

## Introduction

Recently, the gut microbiota has been implicated in somatic pain. In humans, an abnormal composition of the gut microbiota could be associated with chronic post-operative, osteoarthritic pain and fibromyalgia^13–16^. In rodents, animals devoid of gut microbiota (germ free, GF) or in which the microbiota is depleted by antibiotic administration, develop less sensory hypersensitivity in response to tissue inflammation and to peripheral neuropathies induced by chemotherapy, diabetes and injury of the sciatic nerve^8–11^. The reported underlying mechanisms involve the control that the gut microbiota exerts on immune cells. In response to tissue injury, GF mice or mice depleted of gut microbiota, exhibit decreased local production of pro-inflammatory cytokines, decreased macrophage expansion in the DRG, reduced spinal microglial activation and enhanced expansion of regulatory T cells in the spinal cord^8–11^. Together, these studies show that signals from gut microbiota are required for the initiation of injury-induced sensory hypersensitivity. However, whether an abnormal gut microbiota is able to promote pain chronicity remains largely under documented. Here, we report that male mice knock-out (KO) for myosin 1a (*Myo1a*)^12^ raised under single genotype (KO-SGH) but not mixed genotype housing (KO-MGH) conditions exhibit a persistent mechanical hypersensitivity in response to inflammatory, post-operative and neuropathic tissue insults. We demonstrate that the acquisition of the vulnerability to injury-induced chronic pain in KO-SGH is achieved through the inheritance of a dysbiotic microbiota. Parental antibiotic treatment completely rescues the chronic pain phenotype, modifies the gut microbiota composition and resolves the neuropathy-induced immune response in the DRG, in KO-SGH male offspring.

## Results

### Myo1a-deficient males raised under single genotype housing conditions exhibit a vulnerability to injury-induced chronic pain

In a previous study, in which we analysed the transcriptional signature of flow-cytometry activated cell-(FAC-) sorted C-Low-Threshold Mechanoreceptors (C-LTMRs), followed by an extensive *in situ* hybridisation validation, we showed that C-LTMRs share a common gene expression profile with other subsets of LTMRs, including Aδ- and Aβ fibers^17^. LTMRs innervate the skin, from where they convey innocuous mechanical stimuli under physiological conditions, and they also participate to injury-induced mechanical hypersensitivity ^18–20^. Within the common LTMRs set of genes, we identified *Myo1a* as being highly enriched in Gαi-Inhibitory Interacting Protein (GINIP)-positive Isolectin Binding 4 (IB4)-negative C-LTMRs, as expected, but also in Tropomyosin receptor kinase B (*TrkB*)-expressing Aδ LTMRs and in large-diameter Aβ-LTMRs expressing the tyrosine kinase Ret (Fig. 1a-c). In DRG, *Myo1a* was detectable at embryonic day 15 (E15) in a small subset of large-diameter neurons and its expression extended at birth to the majority of DRG neurons (Extended Data Fig.1a-b). In adult mice, *Myo1a* was restricted to LTMRs, as it was undetectable within peptidergic TrkA-positive, non-peptidergic IB4-positive and proprioceptive TrkC-expressing neurons (Extended Data Fig. 1c-e). In addition, *Myo1a*-encoding transcripts were undetectable in the adult SC by qRT-PCR, or in brain areas such as hippocampus and cerebellum (Extended Data Fig. 1f).

**Fig. 1.**
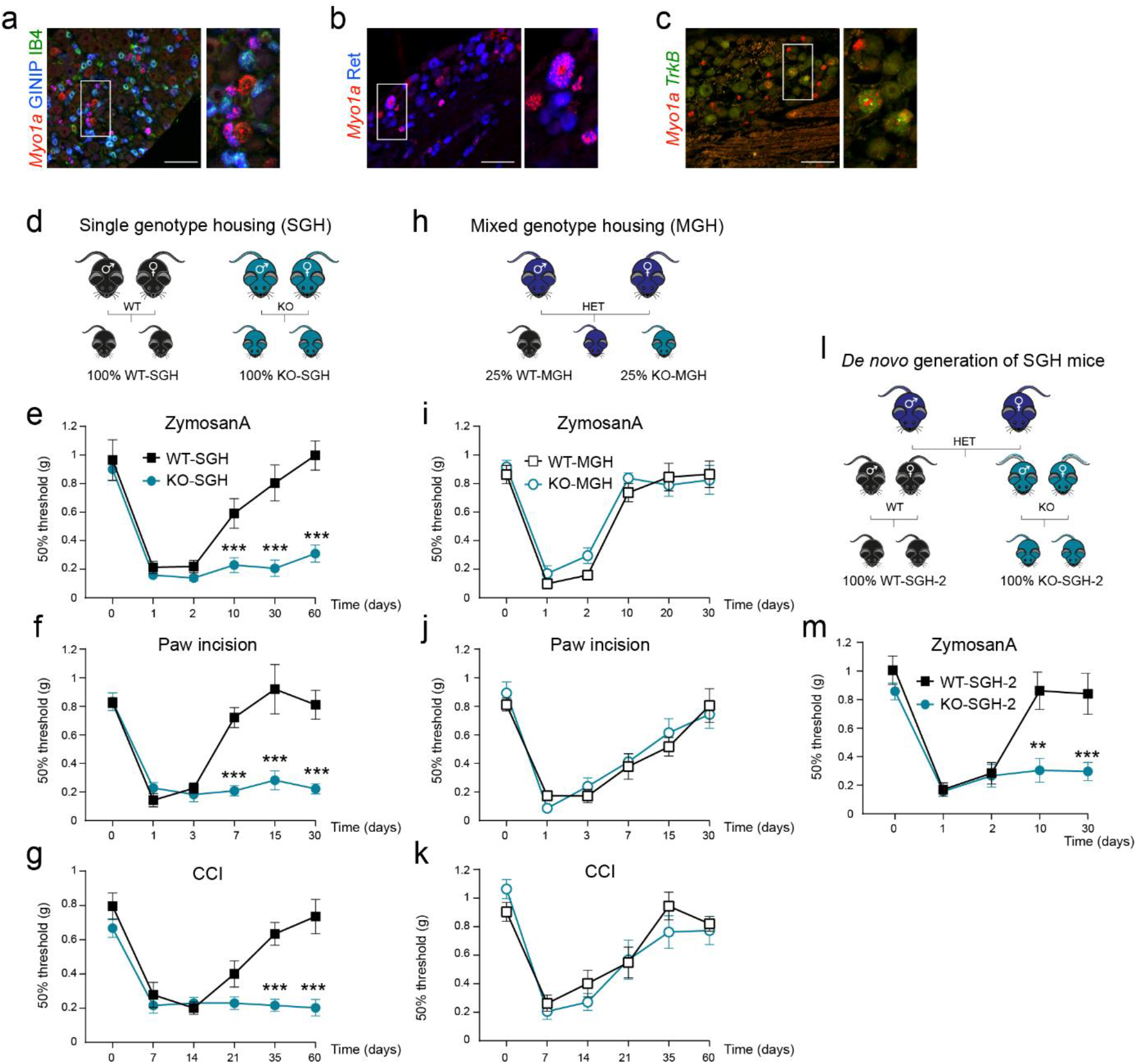
Myo1a-deficient male mice raised under single genotype but not mixed genotype housing conditions exhibit a predisposition to injury-induced mechanical pain. **a-c**, *In situ* hybridization (ISH) showing the expression of *Myo1a* (red) in adult lumbar DRG neurons. **a**, co-labelling with anti-GINIP (blue) and IB4 (green). **b**, double ISH with *TrkB* (green) probe. **c**, co-immunolabelling with anti-Ret (blue). Scale bars 100µm **d**, Schematic representation of breeding and housing of WT and KO mice raised under single genotype housing (SGH) conditions. **e-g** Mechanical thresholds of WT-SGH and KO-SGH males in response to tissue injuries induced by Zymosan A, n=9 per genotype (**e**), paw-incision, n=10 per genotype (**f**) and chronic constriction injury (CCI), n=9 WT-SGH and 10 KO-SGH (**g**). **h**, Schematic representation of breeding and housing of WT and KO mice raised under mixed genotype housing conditions (MGH). **i-k**, Mechanical thresholds of WT-MGH and KO-MGH males in response to tissue injuries induced by Zymosan A, n=9 per genotype (**i**), paw-incision, n=12 WT-SGH and 11 KO-SGH (**j**) and by CCI, n=8 per genotype (**k**). **l**, Schematic representation of the experimental design allowing the de novo generation of WT-SGH-2 and KO-SGH-2 mice from MGH founders. **m**, Mechanical thresholds of WT-SGH-2 (n=11) and KO-SGH-2 (n=14) males in response to Zymosan A. Data are presented as mean ± SEM for each group. ***P<0.01, Two-Way Repeated Measures ANOVA followed by Bonerroni’s multiple comparisons test.

MYO1A caught our attention for three main reasons. First, MYO1A belongs to a large family of atypical type 1 myosins that bridge the cellular membrane to the actin cytoskeleton^21^. Second, MYO1A is highly expressed in small intestinal enterocytes where it plays an important role in microvilli structural organization^12^. Third, MYO1A is closely related to MYO1C which is critical for inner ear hair cell mechano-transduction^22^. On this basis, we sought to investigate the function of MYO1A in somato-sensory biology, by analyzing *Myo1a* Knock-Out (KO) mice^12^. As these mice were available as homozygous breeders and because they were on a pure C57Bl/6J background^23^, we initiated our study by comparing *Myo1a*-KO mice to control C57Bl/6J wild type (WT) mice, which were generated from homozygous crosses. Under these conditions, *Myo1a-*KO and WT mice evolved in single-genotype housing conditions (hereafter respectively called KO-SGH and WT-SGH) (Fig. 1d). Global analysis of KO-SGH mice showed that MYO1A is dispensable for the survival, molecular maturation and target innervation of PSNs, including those that normally express *Myo1a* (Extended Data Fig. 2). Additionally, KO-SGH and WT-SGH mice had similar behaviour in the open filed and rotarod tests (Extended Data Fig. 3a-c). They exhibited no difference in their ability to sense noxious heat and cold stimuli, as shown by their responsiveness in the hot plate test and in the acetone test, respectively (Extended Data Fig. 3d-e).

Because *Myo1a* is expressed in LTMRs, we tested the mechanical sensitivity of KO-SGH and WT-SGH male mice at steady-state and at different time-points following tissue injury elicited by intra-plantar administration of Zymosan A^24^, paw incision surgery^25^ and chronic constriction injury (CCI) of the sciatic nerve^26^. By using the up-and-down method of the Von Frey’s test, we showed that male mice of both genotypes had similar baseline mechanical thresholds and developed significant and similar mechanical hypersensitivity in all three paradigms (Fig. 1e-g). However, in contrast to WT-SGH males in which the injury-induced mechanical hypersensitivity progressively resolved over time, KO-SGH males exhibited persistent mechanical hypersensitivity in all three models of injury-induced pain (Fig. 1e-g). Notably, in the CCI model, while WT-SGH males recovered their baseline mechanical threshold 45 days post-injury, KO-SGH males remained hypersensitive up to 180 days (6 months) post-CCI (Extended Data Fig. 4a). Interestingly, and in line with the selective expression of Myo1a in LTMRs, paw incision-induced thermal hyperalgesia was similar between KO-SGH and WT-SGH mice (Extended Data Fig. 4b). Furthermore, in contrast to KO-SGH males, KO-SGH females did not exhibit persistent mechanical hypersensitivity in response to Zymosan A-, paw incision- and CCI-induced injuries (Extended Data Fig. 5a-c). These data demonstrate that KO-SGH males exhibit a vulnerability to injury-induced chronic pain. They further show that the loss of MYO1A facilitates the transition from acute to chronic injury-induced pain in a sex-dependent manner. Knowing the importance of the interaction between the host and the microbial genomes in determining a given phenotype^27^, we asked whether MYO1A loss-of-function (LOF) *per se* was sufficient to explain our observed phenotype. Towards this aim, we generated WT and KO littermates, evolving in mixed genotype housing conditions (hereafter called WT-MGH and KO-MGH respectively) (Fig. 1h). Under these housing conditions, KO-MGH mice exhibited the same mechanical thresholds as their WT-MGH littermates in response to intra-plantar Zymosan A injection, paw-incision surgery and CCI (Fig. 1i-k). Both groups of mice developed similar injury-induced mechanical hypersensitivity and, in sharp contrast with KO-SGH males, both KO-MGH and WT-MGH exhibited a full recovery of the mechanical sensitivity from tissue injury (Fig. 1i-k). From these data, we conclude that MYO1A loss of function does not play a dominant role in the establishment of the chronic pain phenotype. However, these results strongly suggest that environmental factors may contribute to promoting injury-induced pain chronicity in KO-SGH male mice. Consistent with this hypothesis, *de novo* generation of KO-SGH and WT-SGH second generation male mice from MGH-KO and MGH-WT founders, recapitulates the chronic pain phenotype following intra-plantar injection of Zymosan-A (Fig. 1l-m). Together these data demonstrate that gene-environment interactions are required to trigger the vulnerability to injury-induced chronic pain in myosin 1a-deficient male mice.

### KO-SGH but not KO-MGH male mice exhibit strong alteration in the gut microbiota composition

Knowing that the absence of MYO1A in the gut leads to impaired structure and function of intestinal epithelial cells^12,28^, we hypothesised that the predisposition of KO-SGH males to develop chronic mechanical pain is likely due to an altered gut microbiota. To test this hypothesis, we first performed 16S rRNA gene sequencing from faecal samples of WT and KO male mice raised under SGH and MGH conditions. Principal Component Analysis (PCA) of diversity indices and Linear Discriminant Analysis (LDA) for effect size (LEfSe) revealed a shift in gut microbiota composition of KO-SGH males compared to the WT-SGH counterparts (Fig. 2a-e). This shift was due to an increase in the abundance of rare taxa in KO-SGH, as shown by the Chao-1 index, though the overall diversity was unchanged between groups. KO-SGH male mice had a higher relative abundance of *Olsenella, Parabacteroides, Alistipes and Erysipelatoclostridium* (Fig. 2d-e), previously associated with a disease state such as periodontitis for *Olsenellea*^29^, neurodegenerative disorders and/or depression for *Parabacteroides* and *Alistipes*^30^ and high fat diet for *Erysipelatoclostridium*^31^. In contrast, we found no significant difference in the gut microbiota composition of KO-MGH males as compared to their WT-MGH littermates (Fig. 2f-h). This is in line with previous studies reporting a normalisation of gut microbiota between genotypes, when mice are raised as littermates^32^.

**Fig. 2.**
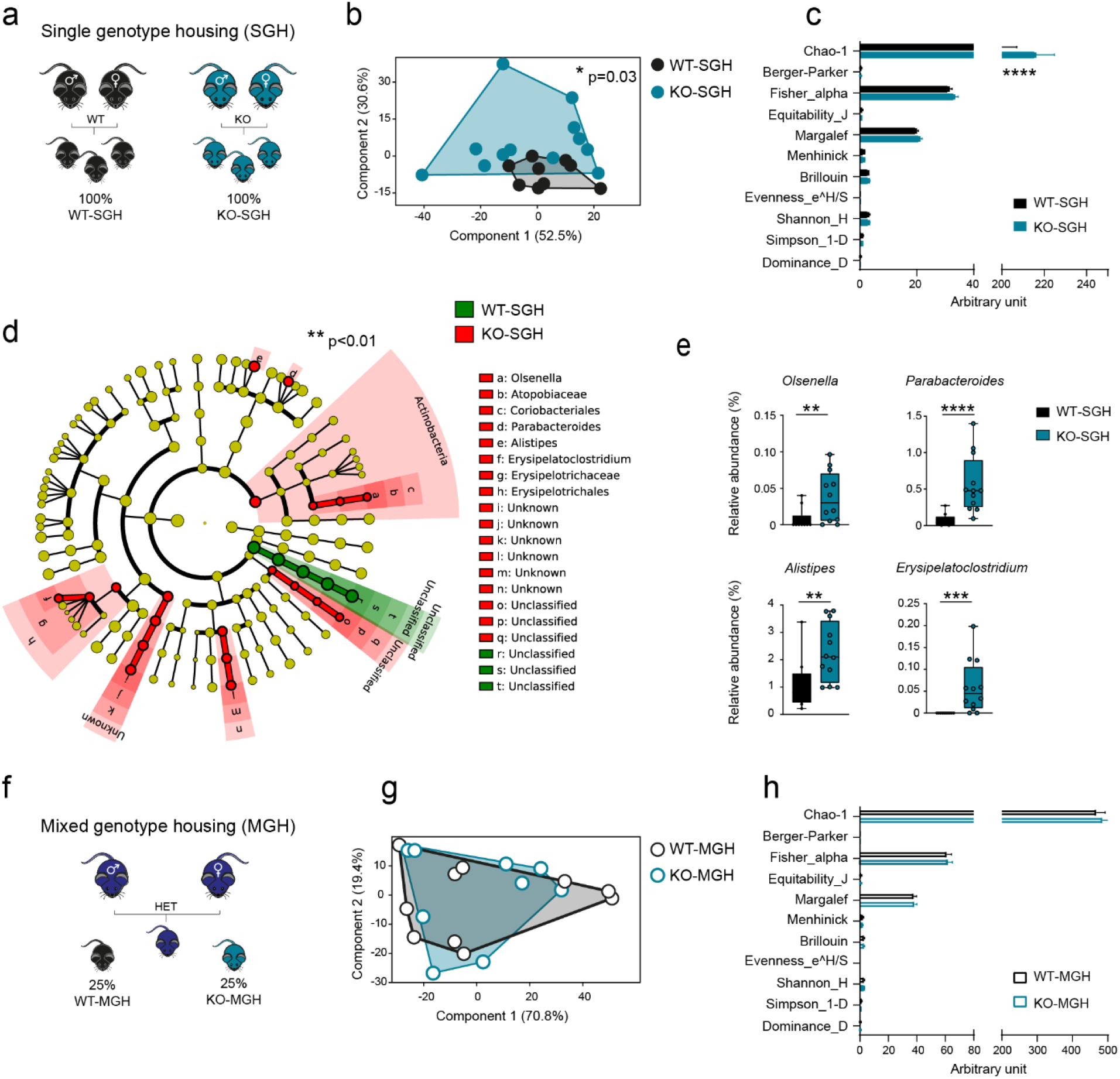
KO-SGH but not KO-MGH males exhibit alterations in the gut microbiota composition. **a**, Schematic representation of breeding and housing of WT-SGH and KO-SGH mice. **b**, Principal component analysis (PCA) of gut microbiota of WT-SGH (n=10) and KO-SGH (n=12) males, \**P*<0.05, 1-way PERMANOVA with Bonferroni correction. **c**, Diversity indices of gut microbiota of WT-SGH and KO-SGH males, \*\*\*\**P*<0.0001, 2-way ANOVA followed by a 2-stage linear step-up procedure of Benjamini, Krieger and Yekutieli to correct for multiple comparisons by controlling the False Discovery Rate (< 0.05). **d**, Cladogram showing bacterial taxa significantly higher in WT-SGH and KO-SGH gut microbiota (the cladogram shows taxonomic levels represented by rings with phyla at the innermost and genera at the outermost ring and each circle is a bacterial member within that level). **e**, Relative abundance of *Olsenella, Parabacteroides, Alistipes* and *Erysipelatoclostridium* in WT-SGH and KO-SGH gut microbiota, \*\**P*<0.01, \*\*\**P*<0.001, \*\*\*\**P*<0.0001, Mann Whitney U test. **f**, Schematic representation of breeding and housing of WT-MGH and KO-MGH mice. **g**, Principal component analysis (PCA) of gut microbiota of WT-MGH and KO-MGH males. **h**, Diversity indices of gut microbiota of WT-MGH and KO-MGH males.

As gut microbiota is known to be vertically transmitted via maternal inoculation ^27,32^, our data strongly suggest that the establishment of the dysbiosis in KO-SGH males results from the inheritance of an altered microbiota. In line with this hypothesis, we found that KO-SGH females also exhibited alterations in their gut microbiota composition as compared to WT-SGH females, despite being protected from injury-induced chronic pain (Extended Data Fig. 5d-f). This shift was due to increased abundance in rare taxa, such as *Olsenella, Parabacteroides distasonis* and *Canidiatus Athromitus* (Extended Data Fig. 5f). Altogether our data underscore the potential of MYO1A loss of function to shape the composition of the gut microbiota and to impact on the susceptibility to chronic pain of male mutants.

### Antibiotic treatment of KO-SGH parents rescues male offspring from chronic pain and reduces the CCI-induced immune response in the DRG

Maternal transmission of gut microbiota to the offspring occurs at birth and the early post-natal period is critical for determining its composition^33^. Our results highlight a strong link between the inheritance of a dysbiotic gut microbiota and the vulnerability of KO-SGH male mice to develop injury-induced chronic pain. To test this, we thus decided to deplete the gut microbiota of KO-SGH parents and to analyse the underlying consequences on their male offspring. Such experimental design also considers possible paternal horizontal transmission, as in our animal facility, both parents are housed with their pups. KO-SGH males and females of six weeks of age were treated with a cocktail of large spectrum antibiotics (ABX) for three weeks^9^, whereupon the treatment was stopped and ABX-KO-SGH breeding pairs were formed (Fig. 3a). We analysed the response of adult male offspring (hereafter named pABX-KO-SGH) to paw incision and CCI as well as their gut microbiota composition (Fig. 3a-c). In both pain models, pABX-KO-SGH male mice were compared to KO-SGH and WT-SGH. While all three groups developed similar paw incision- and CCI-induced mechanical hypersensitivity (Fig. 3b-c), KO-SGH mice remained hypersensitive at day 30 in the paw incision model and at day 60 in the CCI model. Importantly, in both pain models, pABX-KO-SGH male mice exhibited the same kinetics of recovery as the WT-SGH mice, further supporting the idea that the inherited dysbiotic microbiota in KO-SGH males is at the origin of their predisposition to injury-induced pain (Fig. 3b-c). Consistently, following 16S rRNA gene sequencing analyses of faecal samples, PCA and LEfSe revealed a significant shift in the gut microbiota composition of pABX-KO-SGH males compared to SGH-KO males (Fig. 3d-f). In particular, pABX-KO-SGH mice had a statistically significant lower relative abundance of bacteria such as *Prevotella, Alistipes* and *Rikenellaceae* and higher relative abundance of bacteria from class *Bacilli*, such as *Lactobacillus*, including *Lactobacillus animalis*, a probiotic species with beneficial properties^34^(Fig. 3e-f).

**Fig. 3.**
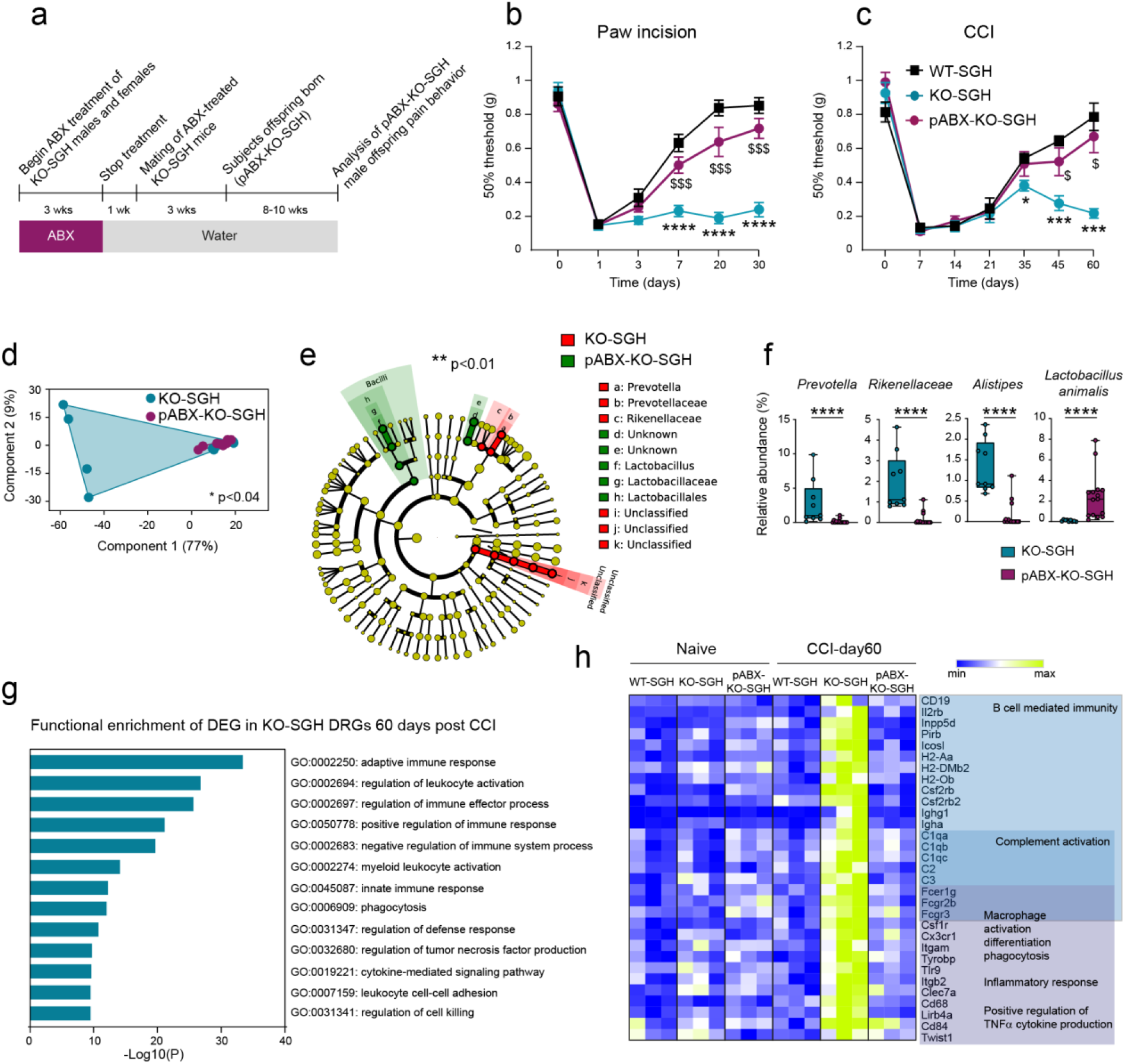
Antibiotic treatment of KO-SGH parents rescues the male offspring from chronic pain and reduces CCI-induced immune response in the DRG. **a**, Experimental design for antibiotic administration to KO-SGH males and females before breeding and subsequent analysis of pABX-KO-SGH male offspring. **b-c**, Mechanical thresholds of indicated mice following paw incision, n=19 WT-SGH, 17 KO-SGH and 21 pABX-KO-SGH and CCI, n=7 WT-SGH, 8 KO-SGH and 7 pABX-KO-SGH (**c**). Mixed effect analysis using Restricted Maximum Likelihood Model and Tukey’s post-hoc test; KO-SGH vs WT-SGH: \**P*<0.05, \*\*\**P*<0.001, \*\*\*\**P*<0.0001, KO-SGH vs pABX-KO-SGH: $*P*<0.05, $$$*P*<0.001, WT-SGH vs pABX-KO-SGH: non-significant. **d**, Principal component analysis (PCA) of gut microbiota of KO-SGH (n=9) and pABX-KO-SGH males (n=15), \**P*<0.05, 1-way PERMANOVA with Bonferroni correction. **e**, Cladogram showing bacterial taxa significantly higher in KO-SGH and pABX-KO-SGH gut microbiota. **f**, Relative abundance of *Prevotella, Rikenellaceae, Alistipes* and *Lactobacillus animalis* in KO-SGH and pABX-KO-SGH gut microbiota, \*\*\**P*<0.001, \*\*\*\**P*<0.0001, Mann Whitney U test. **g**, Gene Ontology (GO) terms enriched in the sets of up-regulated genes in KO-SGH DRGs at 60 days post-CCI. **h**, Heat-map representing the expression levels (Transcripts Per Million) of indicated genes in naïve and CCI-day 60 DRGs from WT-SGH (n=3), KO-SGH (n=3) and pABX-KO-SGH (n=3) males. Shows the biological processes in which these genes are involved, as determined by GO analysis.

To understand the mechanisms underlying the rescue of CCI-induced chronic pain in pABX-KO-SGH mice, we performed RNA deep sequencing of ipsilateral DRG of adult WT-SGH, KO-SGH and pABX-KO-SGH males under naïve conditions and at day 60 post-CCI. Analysis of CCI versus naïve differentially expressed genes (DEG) within the same group and subsequent functional analysis, highlighted differences in the pathways recruited in response to CCI between WT-SGH, KO-SGH and pABX-KO-SGH (Extended Data Fig. 6 and Supplemental Tables 1-3). Since at day 60 post-CCI WT-SGH and pABX-SGH-KO males fully recovered normal mechano-sensitivity while KO-SGH males remained hypersensitive (see Fig. 3c), we sought to identify the sets of preferentially up- or down-regulated DEG only in the KO-SGH CCI-day60 condition. These particular groups of DEG likely reflect those that are associated with pain persistency in KO-SGH males and which are sensitive to regulation by gut-microbiota derived signals. Using, multiplex comparison combined with K-means clustering, we identified a set of 137 DEG up-regulated preferentially in KO-SGH post-CCI (Extended Data Fig. 7 and Supplemental Tables 4-6). Metascape functional analyses^35^ revealed a striking enrichment in genes involved in the immune response (Fig. 3g-h). These included genes involved in adaptive immune response, inflammatory response and positive regulation of cytokine production, demonstrating that several immune effector functions are enhanced in the DRGs of KO-SGH in response to CCI. The set of genes associated with adaptive immune response contained transcripts representative of B-lymphocytes activation (such as cluster of differentiation *Cd19*, and genes belonging to the Major Histocompatibility Complex II: *H2-Aa, H2-Eb1, H2-Oa, H2-Ob*), of antibody production (immunoglobulin heavy chains G1 and A: *Ighg1, Igha*) and of activation of downstream effector functions such as complement recruitment (*C1qa, C1qb, C1qc, C3, C4b, Itgam, Itgb2*) and engagement of activating and inhibitory Fc receptors of effector cells (*Fcer1g, Fcgr3* and *Fcgr2b*). The up-regulation of these sets of genes occurred concomitantly with the up-regulation of genes involved in the inflammatory response and reminiscent of innate immune cells’ recruitment and/or activation, such as myeloid-lineage cells, including monocyte/macrophages (Mo/MΦ). This set contained the Class II transcriptional Activator *Ciita* that enhances the transcription of MHC II-encoding genes, the CD11b-encoding gene *Itgam*, widely expressed on myeloid cells, the Colony Stimulating Factor Receptor 1 *Csf1r*, the chemokine receptor *Cx3cr1* and *Cd68*, reminiscent of Mo/MΦ recruitment, survival and activation, the Pathogen Recognition Receptors *Clec7a*, highly expressed in phagocytic MΦ and in pathogenic microglia^36,37^, including in neuropathic-pain promoting microglia^38^ and the DAP-12 encoding gene *Tyrobp*, also up-regulated in pathogenic microglia and necessary for the induction of neuropathic pain^6,39^.

Overall, these data demonstrate that dybiotic microbiota inherited from KO-SGH parents promotes pain chronicity in KO-SGH male offspring, likely by influencing the immune response in the DRG, following nerve-injury.

## Discussion

In this study, we demonstrate that gut microbiota composition critically determines the persistency of injury-induced chronic mechanical pain, in Myosin 1a knock-out male mice. To our knowledge, this is the first demonstration establishing a causal relationship between a dysbiotic gut microbiota and the predisposition to chronic injury-induced mechanical pain, in preclinical mouse models of pain. Indeed, we found that the acquisition of chronic mechanical pain in KO-SGH male depended on the inheritance of a dysbiotic microbiota. Manipulation of gut microbiota in KO-SGH parents by antibiotic administration completely reversed the mechanical pain chronicity induced by paw incision and CCI, in their pABX-KO-SGH male offspring and promoted the expansion of beneficial bacteria such as *Lactobacillus animalis*^34^. Recent studies on the topic have shown that antibiotic-mediated gut microbiota depletion prevents or decreases the on-set of neuropathic-induced sensory hypersensitivity, in WT, non-genetically manipulated mouse models^9–11^. These include chemotherapy-, diabetes- and CCI-induced neuropathic pain models, with some discrepancies existing for the latter model^9–11^. For instance, antibiotic administration prior to the injury was initially reported to prevent the on-set of mechanical and thermal hyperalgesia^10^, while another group has showed that it decreased thermal hyperalgesia but had no effect on mechanical hypersensitivity^11^. Despite these discrepancies, possibly due to differences in the antibiotic administration protocols, these data demonstrate that signals from gut microbiota are required to generate injury-induced sensory hypersensitivity. In addition, in the same study, Ma and colleagues showed that antibiotic-mediated gut microbiota depletion after tissue injury alleviated mechanical hypersensitivity in a model of diabetic neuropathic pain, suggesting that gut microbiota is also involved in pain perpetuation^11^. Consistently, shifts in the gut microbiota and in the resulting metabolite composition in the serum and spinal cord, occur in CCI-induced pain^40^. There is also increasing evidence for a contribution of gut microbiota in post-operative pain, mainly from human clinical studies. For instance, post-operative pain intensity after upper limb surgery was correlated with the composition and the diversity of gut microbiota^41^. Accordingly, alterations in the gut microbiota composition were also found in patients suffering from chronic post-operative pain following breast cancer^13^. Thus, while our results are consistent with the general concept conveyed by these studies, they additionally demonstrate the necessity to consider gene-microbiota interactions when investigating the prediction of injury-induced pain persistency.

Mechanistically, we found that signals from gut microbiota influence the immune compartment in the maintenance of chronic pain. RNA-sequencing analyses revealed that 60 days post-CCI, the DRGs of KO-SGH males but not WT-SGH exhibited the hallmarks of an on-going immune response, which was rescued by parental antibiotic treatment. We found a striking enrichment in genes involved in various aspects of immune response including a “myeloid-macrophage-like signature” defined by *Itgam, Cx3cr1, Csf1r, Cd68, Clec7a1* and *Tyrobp* reminiscent either of a more activated state of the corresponding cell types, or of an increased number of cells expressing those genes. Previous studies have shown that the protection from injury-induced sensory hypersensitivity by antibiotic-mediated microbiota depletion is achieved by decreasing the inflammatory response mediated by DRG macrophages^9^ and spinal microglia^11^ and by promoting the expansion of spinal Foxp3^+^ regulatory T cells^10^. Moreover, DRG macrophages have been shown to expand in response to nerve-injury, to critically contribute to the initiation and the maintenance of chronic neuropathic pain and to respond to gut microbiota signals^6,9^. Yet, tissue-resident macrophages represent a very heterogenous, plastic pool of cells^42^ and the subtilities of DRG macrophages ontogeny and functional diversity remain to be deciphered. Thus, our data are in line with current literature, demonstrating the ability of the gut microbiota to interact with and to modulate the function of the immune system in the pathogenesis of various conditions, including pain^1^.

Our study also showed that female KO-SGH are protected from injury-induced chronic mechanical pain, despite exhibiting a dysbiosis as compared to their WT-SGH counterparts. Sex differences in pain are increasingly acknowledged, with evidence demonstrating that there are inherent male-female differences in pain processing and modulation^43^. One of the most illustrative examples is the sex-dependent recruitment of the immune system in the settlement of injury-induced pain^44,6,45,46^. As inherent male-female differences in the immune system are acknowledged^47^, one may speculate that KO-SGH females escape pain chronicity because they recruit a branch of the immune system that may be less affected by the dysbiosis. Accordingly, sex-specific effects of gut microbiota in models of Alzheimer’s Disease have been previously reported to be mediated by sex-dependent effect on microglia homeostasis^48,49^.

Finally, our study illustrates the limitations of the use of littermates when a phenotype is mainly driven by the gut microbiota. Indeed, we found that KO-SGH non-littermates but not KO-MGH littermates males exhibited a predisposition to chronic injury-induced mechanical pain, that was accompanied by a dysbiotic microbiota. Remarkably, *de novo* generation of KO-SGH males from KO-MGH littermate founders recapitulated the vulnerability to chronic mechanical pain, demonstrating that MYO1A loss of function generates a detrimental environment promoting the vulnerability to chronic mechanical pain. Because of its expression and function in the small intestine ^12,28^, we hypothesise that MYO1A loss of function affects gut microbiota composition.

Littermate controls are considered as the gold standard to unambiguously prove the dominant contribution of a mutation to a given phenotype ^27^. However, when gene-microbiota interactions come into play, the use of littermates *sensum strictum*, i.e WT and mutant mice originating from heterozygous crossings and continuously housed together, may mask potential shifts in gut microbiota composition caused by the mutation and its influence on the emergence of a given phenotype. Indeed, MGH conditions standardise the composition of gut microbiota across genotypes by normalising the maternal transmission and the horizontal transfer between mice of the same cage^32^. Hence, when KO-MGH males inherit the microbiota from their heterozygous mother and are housed with their WT-MGH littermates, they do not develop injury-induced chronic pain. In sharp contrast, when KO-SGH males inherit the microbiota from their mutant mother and are housed only with mutant mice, they develop injury-induced chronic pain and exhibit alterations in their gut microbiota. Thus, our data underscore the potential of MYO1A loss of function to shape the composition of the gut microbiota and impact on the susceptibility to chronic pain of male mutants. This conclusion fits within the current debate on the impact of a genetic deletion on the composition of gut microbiota^50–52^.

In conclusion our study extends the current knowledge on the link between a dysbiotic gut microbiota and the emergence of injury-induced chronic pain and underscores sex-specific underlying mechanisms, that remain to be deciphered.

## Acknowledgments

We thank all the lab members for their involvement in this project and fruitful discussions. This project was founded by the French National Agency of Research grants ANR-CE16-Myochronic awarded to AM and ANR-CE16-Floradoloris awarded to AR. It was also supported by founding from CNRS and Aix-Marseille University awarded to IBDM.

## Author Contribution

A.R performed ISH and immunostaining experiments, fecal DNA extraction for 16S rRNA sequencing, RNA-extraction for RNA-sequencing, antibiotic treatment, analysed the data and made all the figures. Z.A.J performed all the Von Fery’s behavioural tests on WT-MGH and KO-MGH males, analysed the corresponding data, participated to the analysis of 16S rRNA sequencing and generated unpublished data. S.G performed all the surgeries for CCI and paw incision and analysed the mechanical sensitivity of WT-SGH and KO-SGH males and females (Von Frey’s test). A.Ma performed mouse behaviour experiments (openfield and rotarod and unpublished experiments). P.M performed qRT-PCR experiments, participated to fecal DNA and RNA extraction and to the management of mouse colonies. A.J.S analysed the RNA-sequencing data. M.S performed the taxonomic and predicted functional analysis of gut microbiota. G.H, C.S, F.C and S.U participated to unpublished data. AR and AM designed the project and wrote the manuscript. All the authors provided input and corrections to the manuscript.

## Supplemental Information

Contains Extended Data Fig. 1-7 and Supplemental Tables 1-6.

## Materials and Methods

### Mice

Mice were maintained under standard housing conditions (23°C, 40% humidity, 12 h light cycles, and free access to food and water). Special effort was made to minimize the number as well as the stress and suffering of mice used in this study. *Myo1a* KO mice were generated by Tyska et al., 2005^1^ and were backcrossed for at least 10 generation on C57BL/6 background^2^. Control wild-type (WT) C57BL/6 mice were bred in-house. To generate WT and *Myo1a* KO mice under single genotype housing conditions (WT- and KO-SGH), males and females of each genotype were bred together and the offspring was subsequently analyzed. To generate WT and *Myo1a* KO mice under mixed genotype housing conditions, WT and *Myo1a* KO males were first bred together allowing to generate F0 heterozygous males and females, which were further bred together and the resulting F1 WT-MGH and KO-MGH mice were housed together after weaning. For the *de novo* generation of WT- and KO-SGH mice, WT-MGH males were bred with WT-MGH females and KO-MGH males were bred with KO-MGH females, and the offspring was analyzed.

### Ethical Approval

The authors declare that all protocols are in agreement with European Union recommendations for animal experimentation (EU0221).

### Antibiotic administration

A cocktail of 0.5 g/L ampicillin (Sigma), 0.5 g/L neomycin (Sigma), 0.5 g/L metronidazole (Sigma) and 0.25 g/L vancomycin (Sigma) and 3g/L sucralose, dissolved in water was administrated in the drinking bottles of 6-week-old males and females KO-SGH. The mixture was replaced with a fresh one every 2 to 3 days. After 3 weeks of treatment, the antibiotics were removed, whereupon ABX-KO-SGH breeding pairs were generated and the male pABX-KO-SGH offspring were analyzed.

### Histology

To obtain adult Dorsal Root Ganglia (DRGs) specimens for *in situ* hybridization (ISH) and immunostainings, mice were deeply anesthetized with a mix of ketamine/xylazine and then transcardially perfused with an ice-cold solution of paraformaldehyde 4% in 0.1 M phosphate buffer (4% PFA). Tissues were further fixed for 24h in ice-cold 4% PFA. Newborn P0 mice were sacrificed, rapidly washed in ice-cold PBS, eviscerated and fixed for 24h in ice-cold 4% PFA. E15 embryos were collected in ice-cold PBS and fixed for 24h in ice-cold 4% PFA. Adult back hairy skin was excised and fixed for 2h in ice-cold 4% PFA. Specimens were transferred into a 30% (w/v) sucrose solution for cryoprotection before being frozen in OCT mounting medium. 12μm cryosections (DRGs) and 18-20 µm cryosections (SC, skin, E15 and P0) were obtained using a standard cryostat (Leica).

### In situ hybridization

Dioxigenin-labeled *Myo1a* antisense probe and Fluorescein-labeled *TrkB* probe were synthetized using gene-specific PCR primers and cDNA templates from adult mouse DRG and ISH or double ISH was carried as described in Reynders et al., 2015^3^. The primers used for probe synthesis are listed below.

*Myo1a F1: GAAAATACTTCCGGTCAGGTG*

*Myo1a R1: CAAGGGTTCTTCATCTCTGAGT*

*Myo1a F2: TACCAGTGGAAGTGCAAGAAGT*

*Myo1a R2+T7: TAATACGACTCACTATAGGGACACTACGAAGTTCTGCTCCAG*

*TrkB-F1: CTGAGAGGGCCAGTCACTTC*,

*TrkB-R1: CATGGCAGGTCAACAAGCTA*

, *TrkB-F2: CAGTGGGTCTCAGCACAGAA*,

*TrkB-R2+T7: TAATACGACTCACTATAGGGCTAGGACCAGGATGGCTCTG*

### Immunostaining

Immunostainings were done with rat anti-GINIP (1:500, Moqrich laboratory), goat anti-Ret (1:500, R&D Systems), rabbit anti-TrkA (1:1000, generous gift from Dr. L. Reichardt, University of California), goat anti-TrkC (1:500, R&D Systems), rabbit anti-CGRP (1:1000, ImmunoStar), rabbit anti-PKCγ (1:500, Santa Cruz), rabbit anti-PGP9.5 (1:200, Thermo Scientific) and rabbit-anti S100 (1:1000, Darko). IB4 labelling was performed with Alexa Fluor 488-conjugated IB4 from Invitrogen. Slides were mounted with ImmuMount (Thermo Scientific) prior to observation under AxioImager Z1 (Zeiss) fluorescence microscope. Contrast was adjusted using Photoshop software.

For SC and skin innervation, image acquisition was performed using an LSM-780 confocal microscope (Zeiss) and same pinhole aperture, lasers intensities as well as gain parameters were respected between WT and KO specimens.

### Cell counts

Total and subsets of DRG neurons were counted on lumbar L4 DRG from adult WT and *Myo1a* KO mice, as described in Gaillard et al., 2014^4^. Briefly, 12 µm serial sections of L4 DRG were distributed on 6 slides which were subjected to different markers, including the pan-neuronal marker PGP9.5. This approach allowed us to refer all counting to the total number of neurons (PGP9.5). For each genotype, three independent experiments were performed.

### Behavioral tests and statistical analysis

All behavioral analyses were conducted on 8–12 weeks old KO and WT males and females. All experiments were carried at room temperature (∼22°C). Animals were acclimated for one hour to their testing environment prior to all experiments. Experimenters were blind to the genotype of the mice during testing. The number of tested animals is indicated in the figure legends section.

*Openfield, Rotarod* and *Hot plate* tests were carried as described in Gaillard et al^4^.

#### Acetone test

Acetone drop evaporation assay^5^ was used to assess naïve mice’s sensitivity to innocuous skin cooling. A drop of acetone was applied on the left hind paw using a 1 ml syringe.

The duration of flinching/pain like behavior (seconds) was recorded immediately following acetone application for a total period of 2 minutes. The test was repeated twice and the mean duration of flinching/pain behavior was calculated.

#### Heat hypersensitivity

Hind paw heat hypersensitivity was determined prior and at different time-points (see legend Extended Data Fig. 4) after paw incision surgery, using Hargreaves plantar test, as described in Gaillard et al., 2014^4^.

#### Mechanical thresholds

Mechanical thresholds of the plantar surface were determined using Von Frey’s filaments with the up-down method^6^, previously as described^4^ prior to and at several time-points after inflammation, neuropathy and paw-incision surgery, as indicated in the corresponding figure legends.

### Injury-induced pain models

#### Zymosan-A induced inflammation

20µL of a solution containing 0.06 mg Zymosan-A in 0.9% NaCl (weight/vol, Sigma), were injected subcutaneously into the plantar side of the left hindpaw, using a 30G needled syringe^7^.

#### Chronic constriction injury (CCI)

Unilateral peripheral mono-neuropathy was induced in ketamin/xylasin-anesthetized mice by performing two loosely tied ligatures (with about 1mm spacing) around the common sciatic nerve^8^ using monocryl resorbable suture filaments (6-0, Ethicon, Piscataway, NJ, USA). The nerve was constricted to a barely discernable degree, so that circulation through the epineurial vasculature was not interrupted. After surgery, the animals were allowed to recover in a warming chamber, and then they were returned to their home cages.

#### Paw incision

The paw incision pain model was performed as described by Brennan et al. 1999^9^. Briefly, mice were anesthetised with ketamin/xylasin and a 5 mm longitudinal incision of the plantar face of the right hindpaw, starting from 2-5 mm from the proximal edge of the heel was performed. The plantar muscle was then carefully elevated with forceps and incised longitudinally with a blade, while leaving muscle’s origins intact. The wound was closed with one horizontal mattress suture using 6.0 silk monofilament (Ethicon, Piscataway, NJ, USA) and the wound site was covered with betadin ointment. After surgery, the animals were allowed to recover in a warming chamber, and then they were returned to their home cages.

### Taxonomic and predicted functional analysis of gut microbiota

Feces were collected from individual mice in an Eppendorf tube and immediately frozen on dry ice. The collection was performed between 11AM and 1PM on mice from at least 2 different cages and 2 different breedings. Total DNA was extracted from feces as already reported^10^. The 16S rRNA gene V3-V4 regions were targeted by the 357wf-785R primers and analysed by MiSeq at RTLGenomics (Texas, USA). An average of 16564 sequences per mouse was generated for WT-SGH and KO-SGH mice of each sex presented in Fig. 2b-e and Extended Data Fig. 5; an average of 62880 sequences per mouse was generated for WT-MGH and KO-MGH males in Fig. 2g-h; an average of 55455 sequences per mouse was generated for KO-SGH and pABX-KO-SGH males presented in Fig. 3d-f. Cladograms in were drawn by using the Huttenhower Galaxy web application *via* LEfSe^11^. Principal component analyses (PCA) and Operational Taxonomic Unit (OUT)-based diversity indices were determined with the software PAST4.10.

### RNA extraction

Mice were deeply anesthetized with a mix of ketamine/xylazine and transcardially perfused with 5-10 mL RNA Later (Quiagen). L3 to L4 DRGs were rapidly dissected and RNA was extracted by using RNeasy Micro Kit (Quiagen), according to manufacturer’s instructions. For quality control, RNAs were loaded on a RNA PanoChip (Agilent) and processed with 2100 Bioanalyzer system (Agilent technology).

### High-throughput RNA sequencing and analyses

WT-SGH, KO-SGH and pABX-KO-SGH DRG RNAs were extracted in experimental triplicates from individual mice. High quality RNA (RIN >8.5) was used for sequencing. RNA-seq libraries were prepared using the TruSeq RNA Sample Preparation Kit (Illumina). All libraries were validated for concentration and fragment size using Agilent DNA1000 chips. Sequencing was performed on a HiSeq 2000 (Illumina), base calling performed using RTA (Illumina) and quality control performed using FastQC (http://www.bioinformatics.bbsrc.ac.uk/projects/fastqc) and RSeQC^12^. Sequences were uniquely mapped to the mm10 genome using Subread^13^ (C version 1.4.6-p2) using default values. Reads mapping to gene exons (GRCm38.p4 gene assembly) were counted using featureCounts^14^ (C version 1.4.6-p2). Differential gene expression was performed using exon counts from biological replicates using the DESeq2 BioConductor R package^15^, using a 1% false discovery rate (FDR) and log2 Fold Change of 0.5 cutoff. Heat-maps and K-means clustering with one minus Pearson correlation were generated using Morpheus on-line software https://software.broadinstitute.org/morpheus. Functional analysis was performed using Metascape software ^16^. WT-SGH, KO-SGH and pABX-KO-SGH, CCI vs naïve differentially-expressed genes (DEG) are depicted in Supplementary Tables 1, 2 and 3, respectively. KO-SGH versus WT-SGH, pABX-KO-SGH versus KO-SGH and pABX-KO-SGH versus WT-SGH DEG at day 60 post CCI can be found in Supplementary Tables 4, 5, 6, respectively.

### qRT-PCR

RNA obtained from each sample was converted into cDNA using Superscript III Reverse Transcriptase (Invitrogen). Gene expression was assessed by quantitative PCR (qPCR), using qPCR Sybr-Green master mix (ThermoFisher). Samples were run for 40 cycles on a StepOne qPCR apparatus (Applied Biosystems). The relative quantity of transcripts was determined by normalization to *Gapdh* using the standard ΔCt method.

The primers sequences used for qPCR are:

*Myo1a F: CTACGAGCAGCTTCCCATCT*

*Myo1a R: CCACATTTGCCAAAGCATAG*

*Gapdh F: ATGGTGAAGGTCGGTGTGA*

*Gapdh R: AATCTCCACTTTGCCACTGC*

### Statistical analyses

The n numbers of individuals or experimental replicates and the statistical test are indicated in the corresponding figure legend. Statistical analyses were performed with Prism software.

### Data availability

All high throughput RNA-sequencing data have been deposited with the Gene Expression Ombudsman (GEO) and are available under the accession number GSExxxxxxx. All 16S rRNA sequencing data have been deposited on the Sequence Read Archive (SRA) database under the accession number XXXX.

