## Supplementary material for "Gut microbiota promotes pain chronicity in Myosin1A deficient male mice": Supplemetal Information

Extended Data Fig. 1

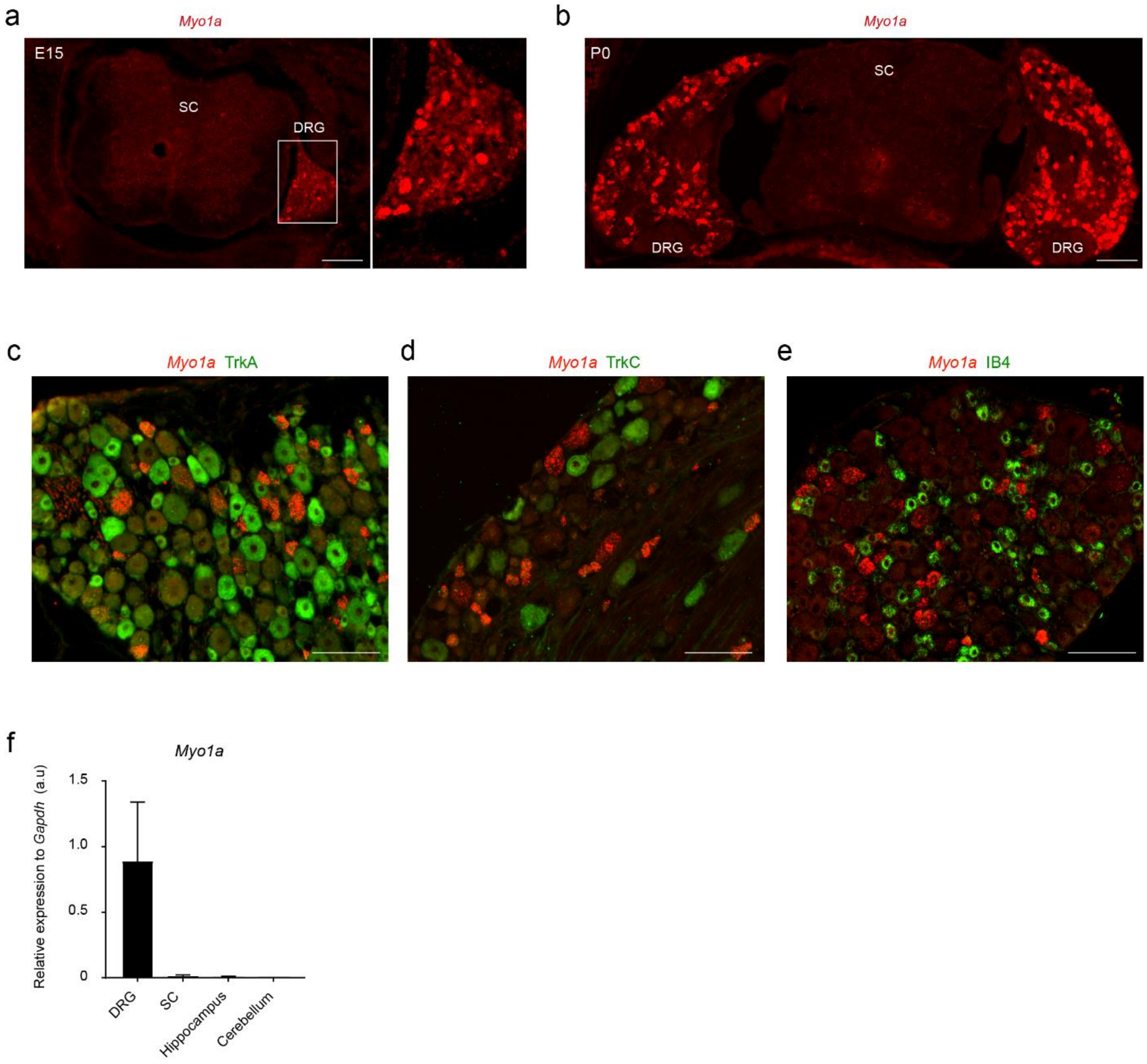


**Extended Data Fig. 1 Expression profile of Myo1a in DRG neurons. a-b**, ISH for *Myo1a* expression on vertebral column cryosections showing DRGs and SC from E15.5 embryos (**a**) and P0 newborns (**b**). Sacle bars:100µm. Data are representative of at least 2 independent experiments. **c-e**: ISH for Myo1a expression in adult lumbar DRGs coupled with anti-TrkA (**c**), anti-TrkC (**d**) immunostainings and IB4 labellind (**e**). Data are representative of at least 3 independent experiments. Scale bars: 50µm. **f**, qRT-PCR showing the expression levels of *Myo1a* in DRG, SC, Hippocampus and Cerebellum relative to *Gapdh*; data are expressed as mean ± SEM of n=2 independent experiments; *a.u*: arbitrary units.

Extended Data Fig. 2

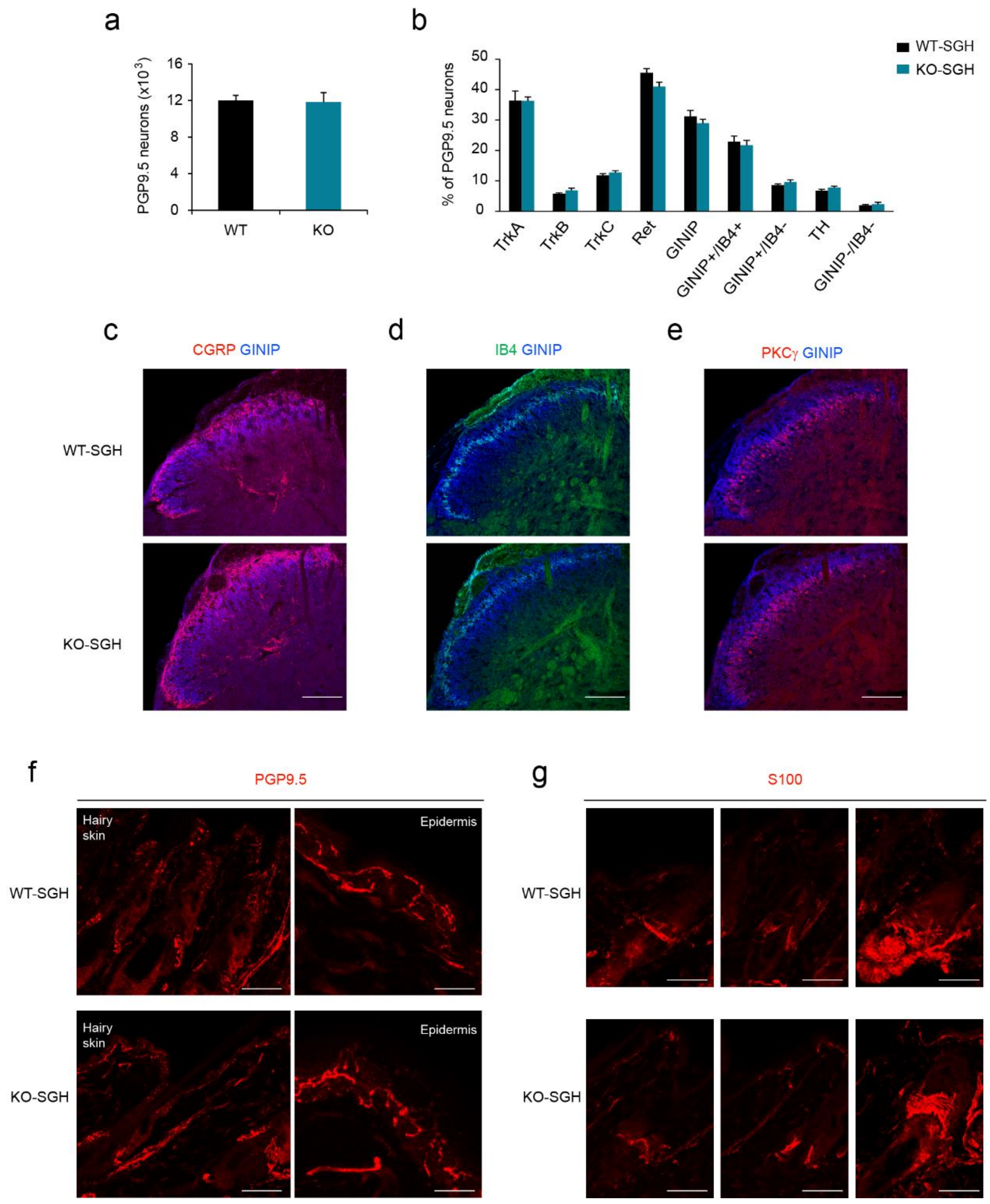


**Extended Data Fig. 2 DRG neurons survival, specification, central and peripheral innervation are not impaired in absence of MYO1A. a**, Cell counts of total PGP9.5-positive neurons in lumbar L4 DRG of WT-SGH and KO-SGH males. **b**, Percentage of indicated neuronal subsets within PGP9.5 neurons determined in the L4 DRG of WT-SGH and KO-SGH males; data are presented as mean ± SEM out of 3 independent experiments (n=3 individuals/genotype). No statistical significance was observed using Mann-Whitney's U test. **c-e**, Immunostaining for GINIP (blue) and CGRP (red) (**c**), IB4 (green) and GINIP (blue) (**d**) and PKCγ (red) and GINIP (blue) (**e**) on lumbar SC cryosections from WT-SGH and KO-SGH males. Scale bars: 100 µm. **f-e** Immunostaining on skin cryosections for PGP9.5 showing hairy (left, scale bars: 50 µm) and glabrous skin (right, scale bars:15 µm) innervation and for S100 showing the innervation of hair follicles (scale bars: 50 µm) in WT-SGH and KO-SGH males. For all immunostaining experiments, data are representative of at least 2 independent experiments.

Extended Data Fig. 3

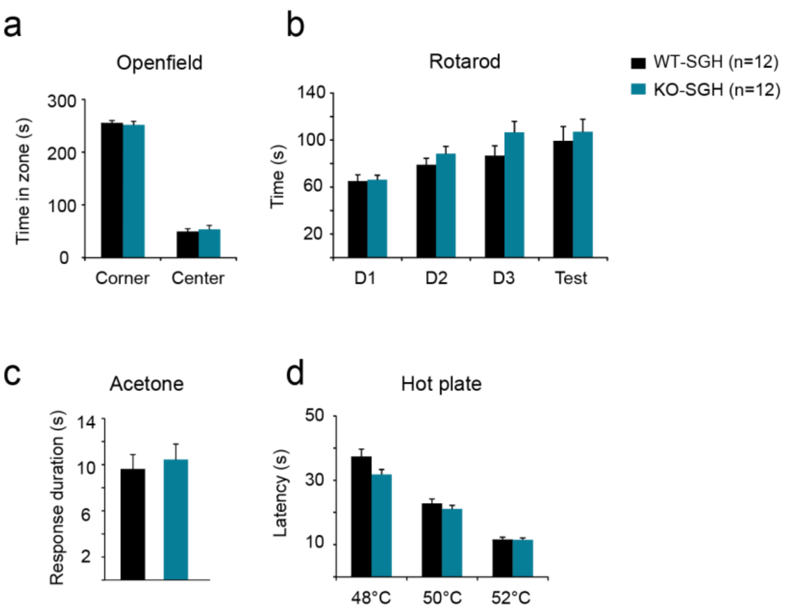


**Extended Data Fig. 3 Naïve KO-SGH males do not exhibit major behavioral deficits. a**, Time spent in center and in corner, in the openfield test. **b**, Time spent on the rotarod apparatus on three consecutive days and during the test day. **c**, Duration of unpleasant response to acetone spray. **d**, Latency before adopting an escape behavior (jump) in the hot-plate teste, at indicated temperatures. All data are presented as mean ± SEM of n=12 WT-SGH and KO-SGH males. No statistical significance was observed using Mann-Whitney’s U test.

Extended Data Fig. 4

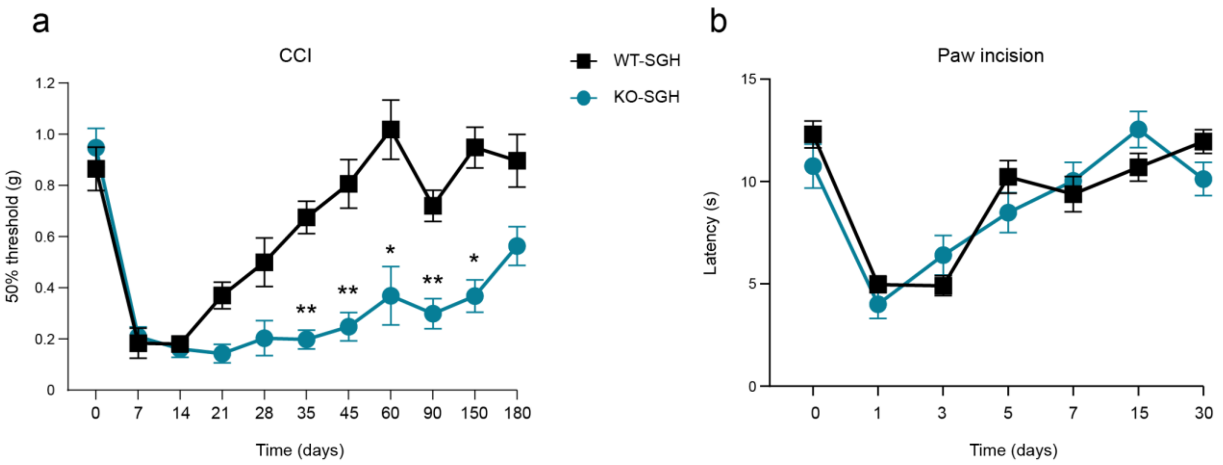


**Extended Data Fig. 4 KO-SGH males exhibit irreversible mechanical hypersensitivity, but not thermal hyperalgesia. a**, Mechanical thresholds of WT-SGH (n=6) and KO-SGH (n=9) males before and at indicated time-points following CCI. *$P$<0.05, **$P$<0.01, Two-Way Repeated Measures ANOVA with Bonferroni's post-hoc test. **b**, Paw withdrawal latency to thermal stimulations in the Hargraves test prior and at different time-points following paw incision, of WT-SGH (n=8) and KO-SGH (n=10) males. No statistical significance between genotypes was observed using Two-Way Repeated Measures ANOVA with Bonferroni's post-hoc test. Data are presented as mean ± SEM.

Extended Data Fig. 5

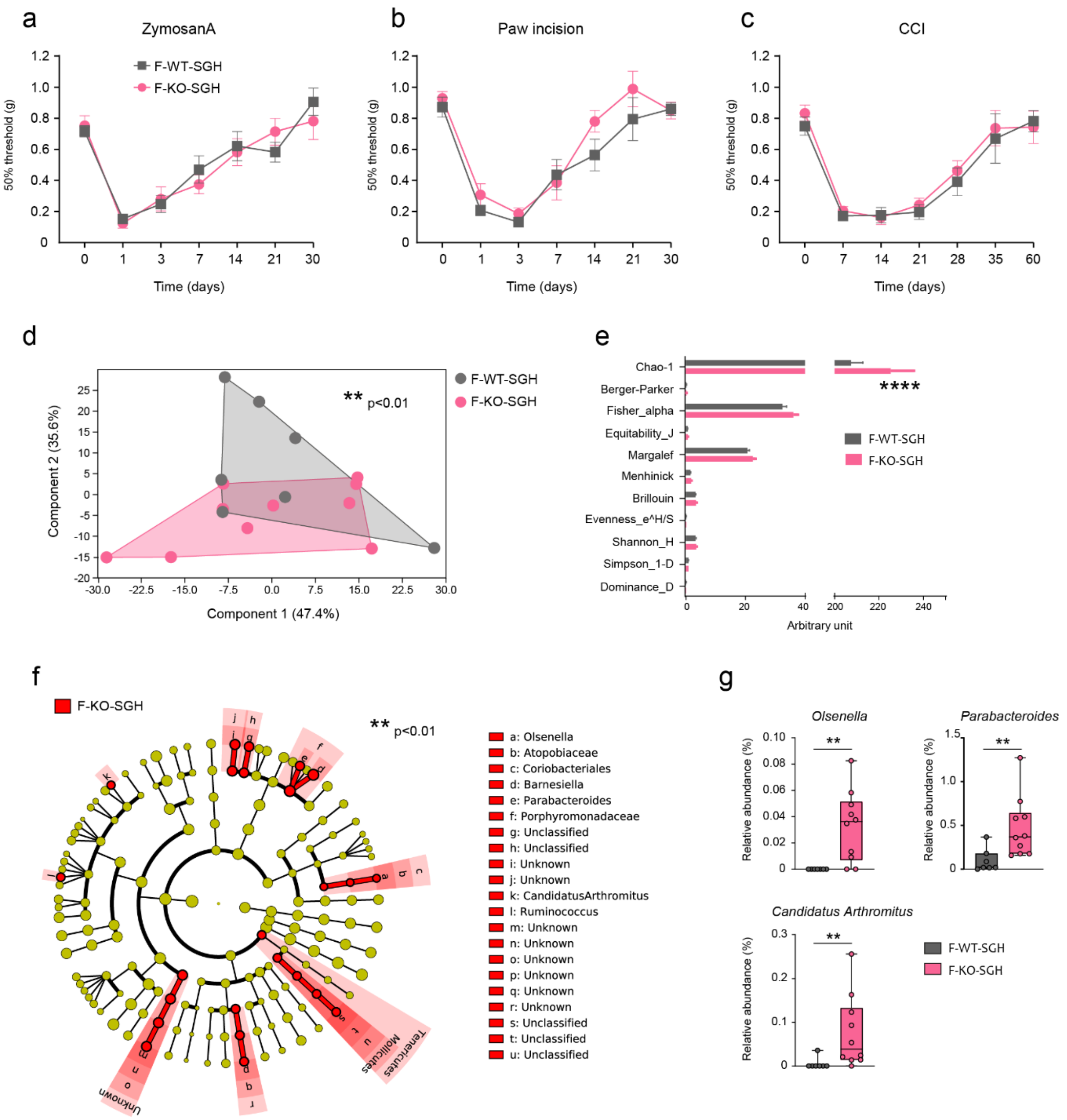


**Extended Data Fig. 5 KO-SGH females do not develop injury-induced chronic pain but exhibit abnormal microbiota composition. a-c** Mechanical thresholds of WT-SGH and KO-SGH females before and at different time points following tissue-injury induced by Zymosan A, n=9 mice per genotype (**a**); paw incision, n=6 mice per genotype (**b**) and CCI, n=7 WT-SGH and 10 KO-SGH (**c**). No statistical significance was observed using Two-Way Repeated Measures ANOVA followed by Bonferroni's post-hoc test. **d**, Principal component analysis (PCA) of gut microbiota of WT-SGH (n=7) and KO-SGH (n=10) females, ***P*<0.01, 1-way PERMANOVA with Bonferroni correction. **e**, Diversity indices of gut microbiota of WT-SGH and KO-SGH females, *****P*<0.0001, 2-way ANOVA followed by a 2-stage linear step-up procedure of Benjamini, Krieger and Yekutieli to correct for multiple comparisons by controlling the False Discovery Rate (< 0.05). **f**, Cladogram showing bacterial taxa significantly higher in female KO-SGH gut microbiota. **e**, Relative abundance of *Olsenella*, *Parabacteroides* and *Candidatus Arthromitus* in female WT-SGH and KO-SGH gut microbiota, ***P*<0.01, Mann Whitney U test.

Extended Data Fig. 6

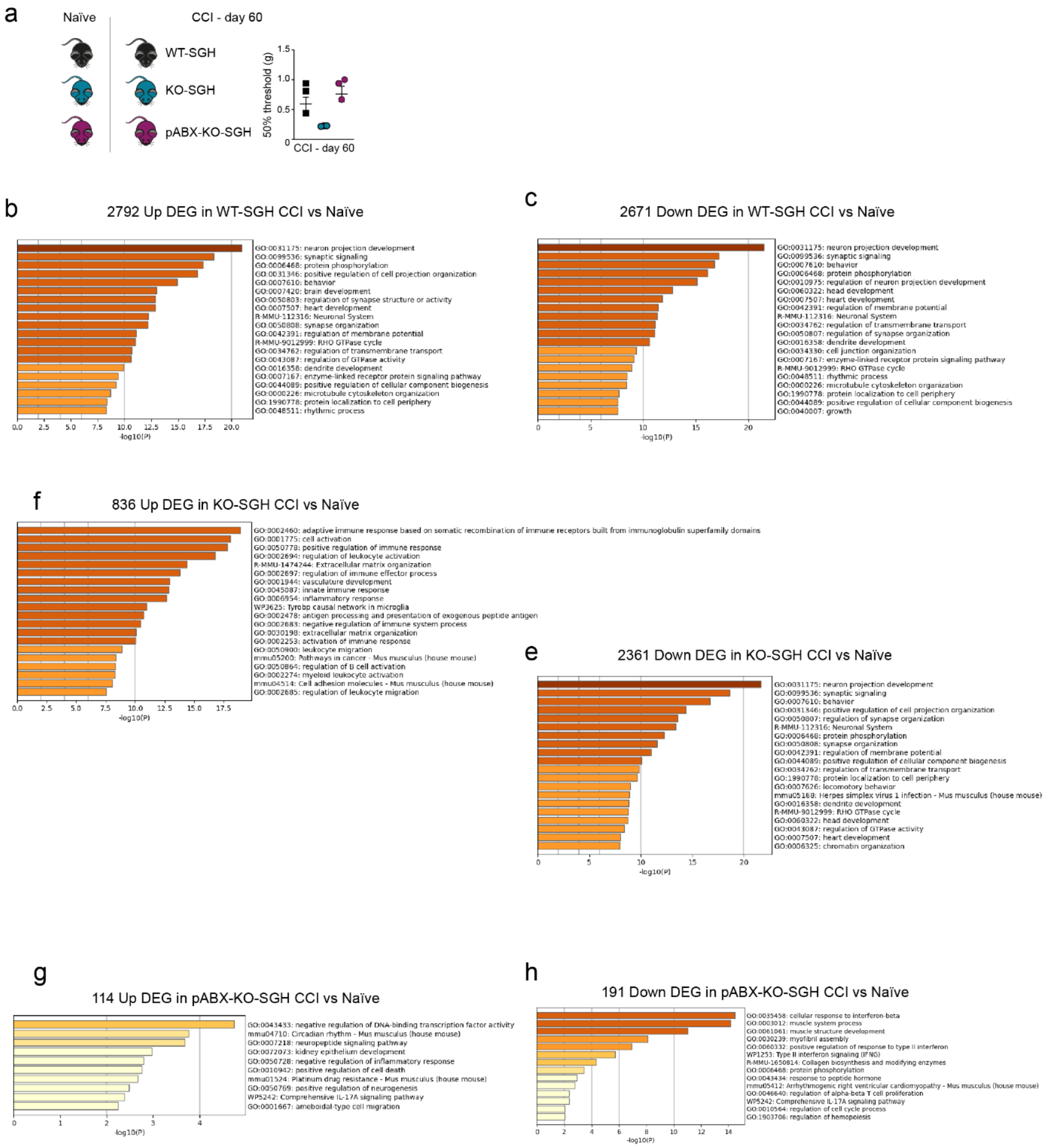


**Extended Data Fig. 6 Different transcriptional programs in the DRGs of WT-SGH, KO-SGH and pABX-KO-SGH males post CCI. a**, Shows the experimental design and the mechanical threshold of the individuals used for RNA-Seq at 60 days post-CCI. **b-h**, Number of Up- and Down-regulated genes in CCI versus naïve conditions and corresponding functional enrichment in WT-SGH males (**b-c**), KO-SGH males (**d-e**) and pABX-KO-SGH males (**g-h**). Differentially expressed genes (DEG) were identified with DESeq2 package36 using a false discovery rate threshold of 0.01 and a cut-off in log2 fold-change (FC) of 0.5.

### Extended Data Fig. 7

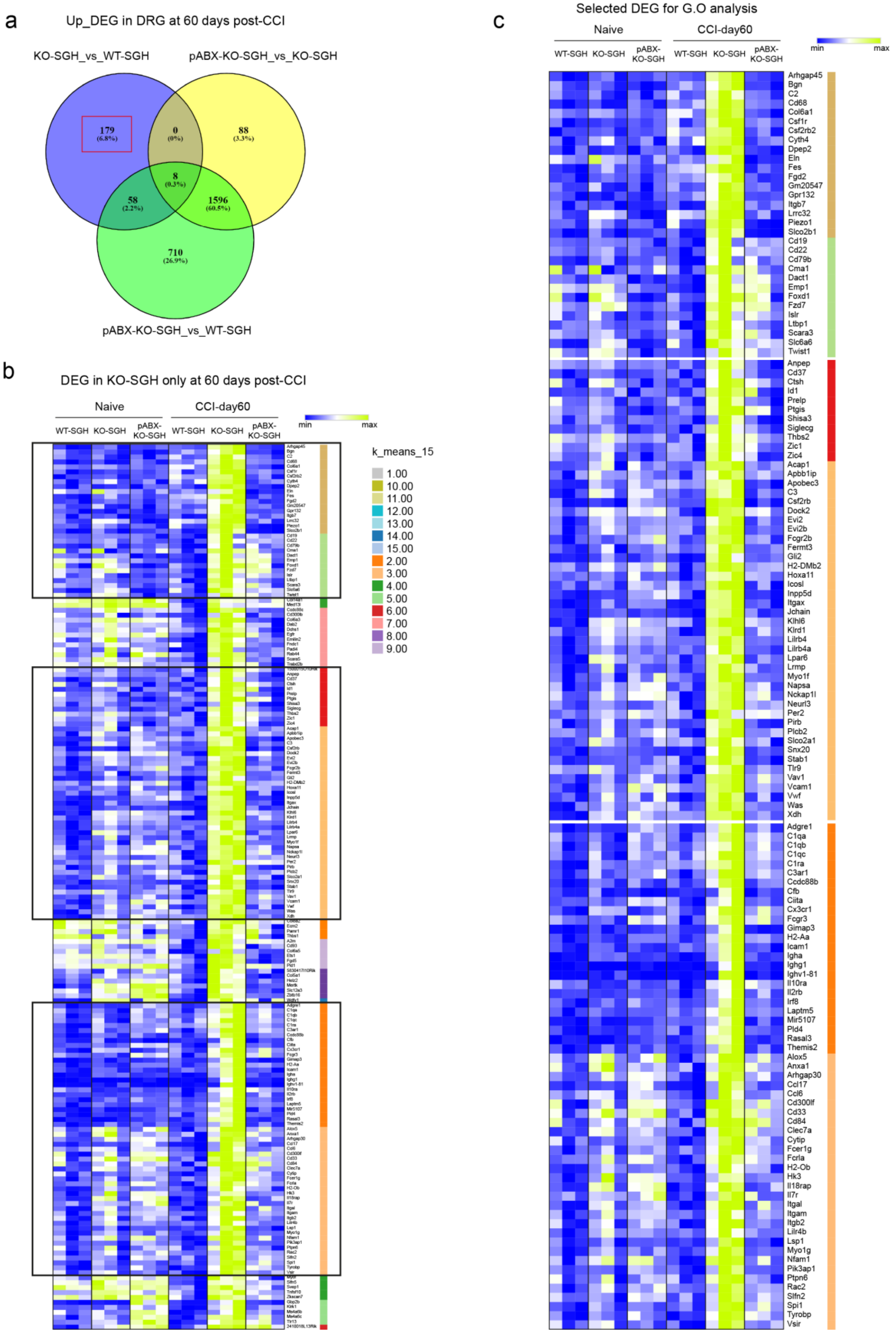

**Extended Data Fig. 7 Selection of sets of genes preferentially expressed in KO-SGH males at 60 days post-CCI. a**, Venn diagram of the up-regulated genes at 60 days post-CCI between KO-SGH and WT-SGH, pABX-KO-SGH and KO-SGH and pABX-KO-SGH and WT-SGH. The genes identified as unique to the KO-SGH condition are circled in red. **b**, Heatmap representation of the expression levels (Transcripts Per Million) and K-mean clustering of the set of genes identified in **a**. **c**, Heatmap representation of the set of genes considered as highly enriched in KO-SGH males at 60 days post-CCI, as compared to the other conditions.
